# AMiGA: software for automated Analysis of Microbial Growth Assays

**DOI:** 10.1101/2020.11.04.369140

**Authors:** Firas S. Midani, James Collins, Robert A. Britton

## Abstract

The analysis of microbial growth is one of the central methods in the field of microbiology. Microbial growth dynamics can be characterized by growth parameters including carrying capacity, exponential growth rate, and growth lag. However, growth assays with clinical isolates, fastidious organisms, or microbes under stress often produce atypical growth shapes that do not follow the classical microbial growth pattern. Here, we introduce the Analysis of Microbial Growth Assays (AMiGA) software which streamlines the analysis of growth curves without any assumptions about their shapes. AMiGA can pool replicates of growth curves and infer summary statistics for biologically meaningful growth parameters. In addition, AMiGA can quantify death phases and characterize diauxic shifts. It can also statistically test for differential growth under distinct experimental conditions. Altogether, AMiGA streamlines the organization, analysis, and visualization of microbial growth assays.

**IMPORTANCE:** Our current understanding of microbial physiology relies on the simple method of measuring microbial populations’ size over time and under different conditions. Many advances have increased the throughput of those assays and enabled the study of non-lab adapted microbes under diverse conditions that widely affect their growth dynamics. Our software provides an all-in-one tool for estimating the growth parameters of microbial cultures and testing for differential growth in a high-throughput and user-friendly fashion without any underlying assumptions about how microbes respond to their growth conditions.

## INTRODUCTION

The study of the growth of microbial cultures has been a basic method of understanding bacterial physiology since the pioneering work of Jacob and Monod and the Copenhagen group in the 1950s-60s (1). Today, there are many automated platforms equipped with multi-well plate readers that can rapidly generate large sets of microbial growth data. Several computational tools have been adopted for the rapid analysis and interpretation of these growth data sets (2–7). However, the growths of clinical isolates, fastidious organisms, or microbes under various stressors often generate curves that do not follow standard logistic or sigmoidal shape. Popular tools using classical mathematical models of growth, such as Logistic or Gompertz equations, struggle to infer kinetic parameters for these microbial cultures. Nonparameteric statistical approaches, including spline fitting or Gaussian Process (GP) regression, are more effective at modelling the growth of atypical microbial cultures and estimating their growth parameters (8–10). Unlike spline fitting, GP regression is robust to outliers and technical variation, can inherently pool replicates, infer derivatives, and estimate growth parameters without cross-validation or bootstrapping (9, 10). User-friendly software to model microbial growth curves with GPs are however lacking which discourages its adoption.

Here, we describe and showcase a new tool for the Analysis of Microbial Growth Assays (AMiGA) without any underlying assumptions about the shape of growth curves. AMiGA models growth curves with GP regression and infers biologically meaningful microbial growth parameters including maximum specific growth rate (i.e. exponential growth rate), lag time, carrying capacity, and area under the curve. Because GPs do not assume any underlying shape for the input observations, we show that AMiGA can quantify adaptation time, death, maximum death rate, and diauxic shifts. Finally, AMiGA can expand growth curves beyond time to include other experimental variables such as nutritional state (e.g. presence of substrate in culture), environmental conditions (e.g. pH status), microbial stressors (e.g. antibiotics), or phylogenetic identities (e.g. genotype). Users can accordingly test for functional differences in growth across distinct experimental conditions (10, 11). These statistical tests are agnostic to the microbial growth parameters but rather detect differences between growth curves across all time measurements. Altogether, AMiGA streamlines various aspects of microbial growth data analysis including quality control, data manipulation, growth curve fitting, and statistical testing.

To illustrate the ease and utility of AMiGA, we demonstrate the inference of microbial growth parameters on several clinical *Clostridiodes difficile* isolates. In particular, we showcase how AMiGA can fit different growth dynamics, characterize diauxic shifts, describe Biolog Phenotype Microarray (PM) plates, and analyze standard growth assays. We also showcase how AMiGA-based testing for differential growth across distinct experimental conditions can extract useful insight from high-throughput growth assays.

## RESULTS

### Implementation

AMiGA is an open-source, cross-platform, Python 3 package. Users interact with AMiGA via the command-line interface. We provide detailed tutorials to demystify the process for users with minimal background in using a command terminal. The main input to AMiGA are raw data files in text format which are often exported by multi-well plate readers. Users can also pass meta-data about each plate or each well using tables saved as text files. AMiGA can recognize Biolog PM plates based on file names and automatically assign wells to carbon, nitrogen, or phosphorous substrates. Users can optionally adjust many of the default parameters for analysis or visualization with a configuration file. AMiGA can analyze multiple data sets in a single batch and it can pool biological and technical replicates before jointly modelling their growth curves.

### Estimating microbial growth parameters

Using observed measurements of optical density (OD), we often want to describe the underlying function of growth often termed the growth curve. AMiGA first transforms OD measurements with a natural logarithm then shifts measurements such that the baseline at the first time point is centered at zero. Using GP regression, AMiGA infers the underlying growth curve which quantifies microbial community size over time and its first-order derivative which quantifies the rate of growth over time (**Fig. 1A and B**). The algorithm then infers biologically meaningful growth parameters by either directly analyzing the growth and its derivative or by sampling from the posterior distribution of the predicted growth model. The sampling approach provides summary statistics for each of the growth parameters in terms of mean, standard deviation, and confidence interval. The estimated microbial growth parameters include classical growth characteristics such as the carrying capacity (K): the maximum growth supported by the environment; area under the curve (AUC): the total growth supported by the environment over observed time; maximum growth rate (r): the maximum specific growth rate or exponential growth rate; doubling time: the time needed during exponential growth to double community size; and lag time: the time delay needed to initiate exponential growth, defined as the intersection of the tangent at maximum growth rate with the axis of time. In addition, AMiGA infers less commonly studied but useful growth characteristics including maximum death rate: the most negative growth rate after reaching carrying capacity; death: the total loss of growth after reaching carrying capacity; and adaptation time: the time interval needed to initiate a positive growth rate. Collectively, these growth parameters describe informative dynamics about microbial physiology under the specified experimental conditions.

**Figure 1.**
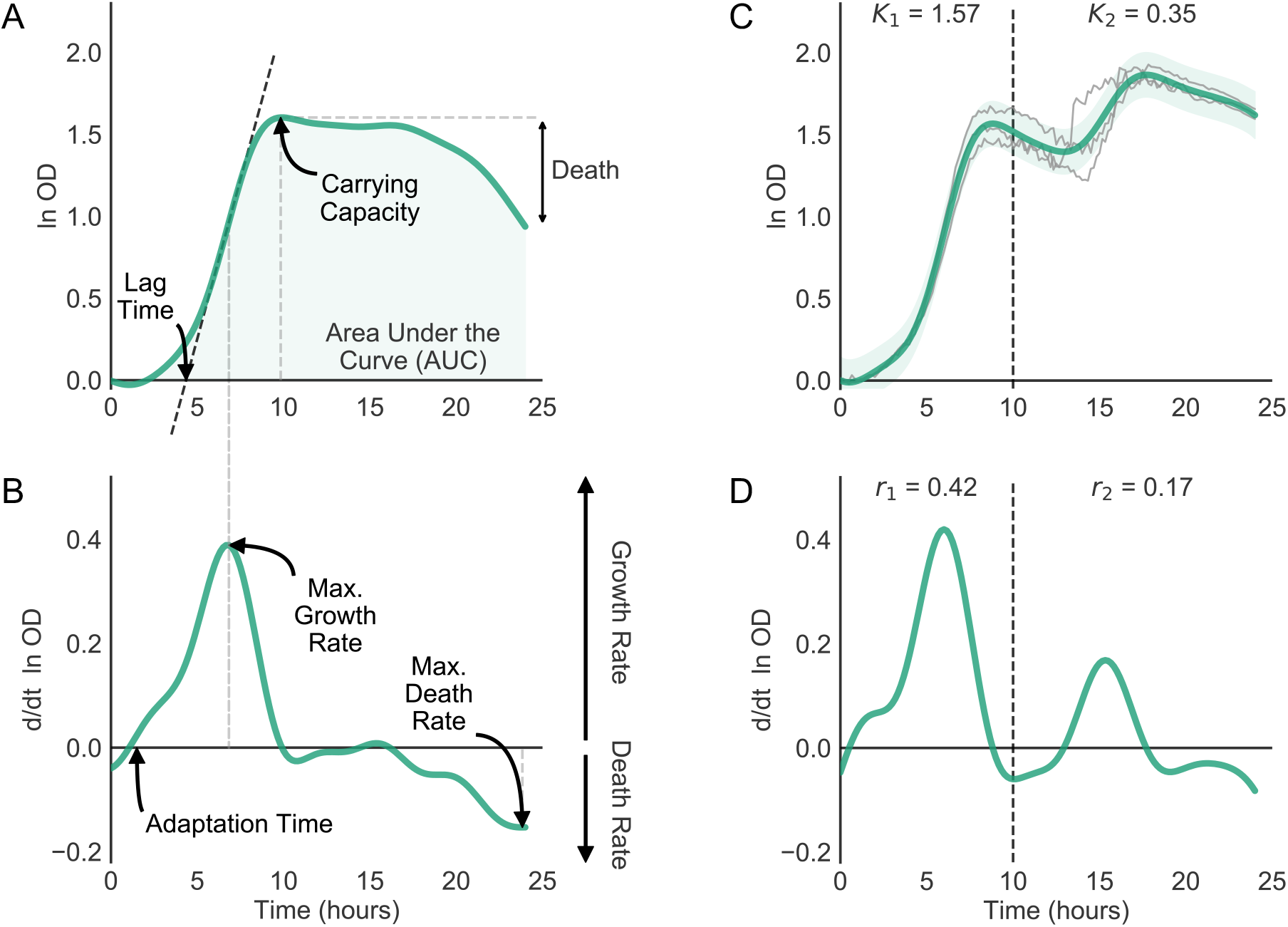
AMiGA infers microbial growth parameters and characterizes diauxic shifts. (**A-B**) AMiGA predicted the mean growth and rate of growth of a ribotype 053 *C. difficile* clinical isolate grown on 20 mM fructose as the sole carbon source using three technical replicates. Model estimates of the log-transformed growth and its derivative can be used to describe the following parameters: carrying capacity (K), area under the curve (AUC), maximum growth rate (r), maximum death rate, lag time, adaptation time, and stationary death. Black dashed oblique line indicates the tangent to the growth curve at maximum exponential growth. Gray dashed vertical lines map growth parameters to their respective time points which are also exported by AMiGA. (**C-D**) AMiGA characterized the diauxic shift of a *C. difficile* clinical isolate of an unknown ribotype grown on minimal media with 20 mM glucose as the sole carbon source using three technical replicates. AMiGA automatically detected two growth phases (separated by dashed vertical line) reaching maximum growth rates at 6.0 and 15.3 hours respectively. Inset text states the estimated carrying capacity, *K*, and maximum growth rate, *r*, for each unique phase. Bold green lines plot the predicted mean of the growth function or its derivative and green bands indicate the predicted 95% confidence interval including measurement noise. Thin gray lines plot actual growth.

### Characterizing diauxic shifts

Diauxie is the biological phenomenon observed when a microbial culture undergoes two phases of growth (1). Diauxic growth often occurs when a microbial culture initially utilizes the most preferred carbon source in its environment but switches to a secondary source once the former is depleted (12). To identify diauxic shifts, we take advantage of a key feature of GPs: the derivative of a Gaussian Process is also a Gaussian Process. Therefore, we can easily infer the first- and second-order derivatives of each growth curve (9). Here, the first-order derivative estimates the growth rate over time, while the second-order derivative estimates the change in the growth rate over time, which is a measure of the acceleration or deceleration of growth. Using the second-order derivative, we identify the inflection points in each growth curve to decompose the curve into sub-components which are potentially unique growth phases. AMiGA then applies a novel iterative process and growth curve thresholds (see **Methods**) to determine if each growth phase is unique or simply an extension of an adjacent growth phase. Here, secondary growth phases are considered unique if they result in a total change in OD of at least 20% of the total change in OD during the primary growth phase (**Fig. S1)**. Because GP regression does not assume any underlying shape of growth, it can capture two or more unique growth phases. It can also detect diauxic shifts with slow transition (**Fig. 1C and D**) or rapid transition into secondary growth as we see for *C. difficile* on minimal media supplemented with sorbitol or ribose (**Fig. 2A and S1**).

**Figure 2.**
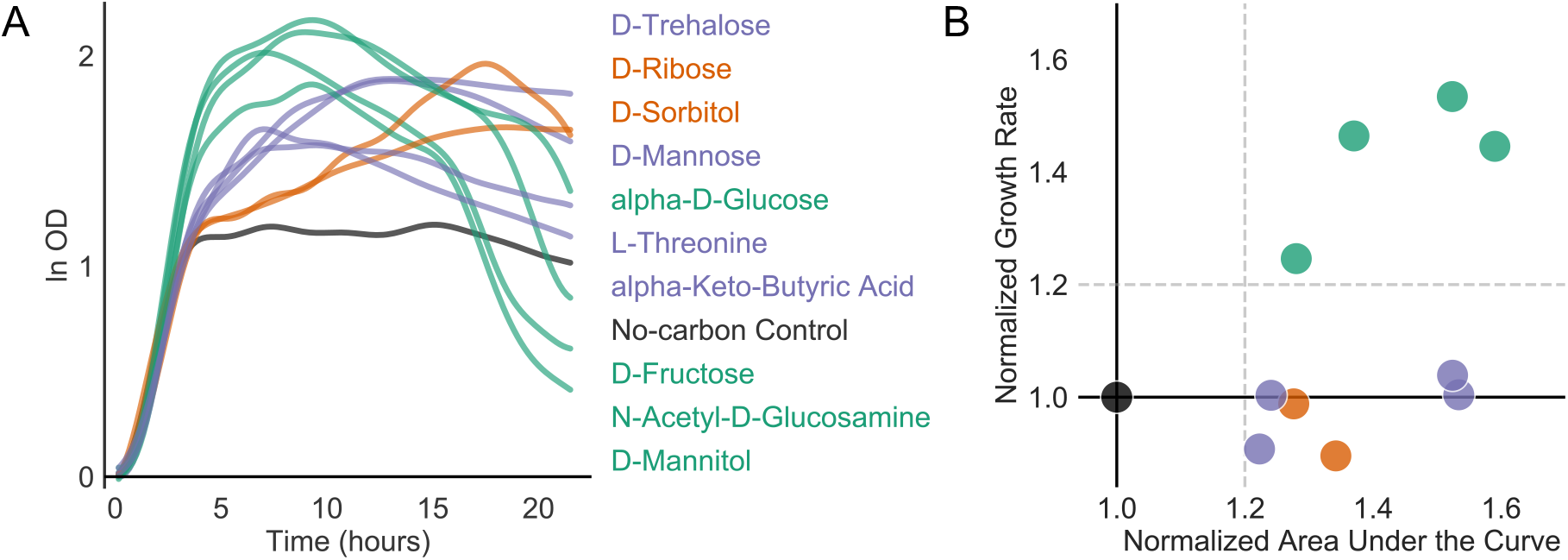
*C. difficile* exhibited growth curves with varying shapes which can be distinguished by growth parameters inferred by AMiGA. (**A-B**) CD2015, a Ribotype 027 *C. difficile* isolate, was profiled with a Biolog Phenotype Microarray (PM1) plate with two technical replicates. Growth curves for each substrate were natural log-transformed, baseline-corrected, then modeled jointly with GP regression by AMiGA. Growth parameters were normalized to the no-carbon control. (**A**) CD2015 grew to a normalized AUC higher than 1.2 on ten substrates. Substrate labels on the right are color-coded and ordered by the final value of their corresponding growth curves. (**B**) On four substrates (green), CD2015 experienced growth rates that are at least 20% higher than growth rate on the no-carbon control. On ribose and sorbitol, *C. difficile* exhibited diauxic shifts (orange). On remaining substrates (purple), *C. difficile* showed logistic or sigmoidal growth and grew at rates comparable to the rate of growth on the no-carbon control (black). Dashed horizontal and vertical lines show arbitrary thresholds for color coding by normalized growth rate and defining positive growth respectively.

### Modelling microbial growth dynamics

*C. difficile* is a gram-positive spore-forming pathogen that has recently become the most common hospital-associated infection in the developed world (13). *C. difficile* is a genetically diverse species and distinct ribotypes are overrepresented in both human outbreaks and animals (14–16). Carbon substrate utilization by clinical *C. difficile* isolates demonstrates phenotypic diversity within and between ribotypes. Here, we profiled CD2015, a ribotype 027 *C. difficile* clinical isolate, on Biolog Phenotype Microarray Plate (PM1) using two technical replicates. All wells in a Biolog PM plate are pre-arrayed with a single carbon source except for the first well, which lacks a carbon source and serves as a no-carbon control.

Using AMiGA, we pooled duplicate growth curves for each substrate, modeled them jointly with GP regression, and inferred microbial growth parameters (**Data S1**). To identify substrates that supported CD2015 growth, parameters were normalized by AMiGA to the no-carbon control well. Predicted growth curves and parameters can be visualized with basic but customizable figures (**Fig. S2 and S3**). We detected significant growth (defined as normalized AUC of at least 1.2) on ten carbon substrates (**Fig. 2**). Because GP regression is a nonparametric approach, AMiGA can fit growth curves with varying shapes (**Fig. 2A**). In minimal defined media supplemented with a single carbon source, CD2015 exhibited multiple growth modalities including rapid growth followed by rapid death (green, Normalized Growth Rate ≥ 1.2), diauxic growth with a slower growth rate in the second phase for a prolonged period of time (orange, Diauxie = True), and logistic or sigmoidal growth (purple and black, Normalized Growth Rate < 1.2) (**Fig. 2B and S4**). As a bacterial generalist, *C. difficile* can colonize different nutritional niches in the gut (17). It is also capable of using amino acids as its sole energy and biomass source (18), which explains its ability to grow on the no-carbon control and may explain its biphasic growth on ribose and sorbitol. *C. difficile* may initially ferment amino acids via the Stickland pathway until they are depleted then transition to growth on sugars present in the environment (18, 19). However, it is less clear why rapid growth on certain monosaccharides was swiftly followed by rapid death.

A parameteric approach for modelling growth would have poorly characterized most of these curves and misrepresented growth dynamics on different substrates. In addition, analysis that eschews fitting growth curves due to their atypical shapes and simply analyzes areas under the curve or carrying capacity would miss difference in growth dynamics. For example, CD2015 showed similar total growth on fructose and trehalose (95% confidence intervals for AUC, in units of ln OD x h, are 35.89-36.55 and 35.93-36.94 respectively) but their growth rates were different (95% confidence intervals are 0.57-0.64 h^−1^ and 0.33-0.48 h^−1^ respectively). Our nonparameteric approach that models different growth shapes is especially useful for high throughput screens for which manual validation of growth curves would be prohibitively laborious.

### Detecting differential growth by comparing growth parameters

In our Biolog phenotyping, we noticed that certain monosaccharides promoted an atypical growth curve characterized by rapid rise followed by a rapid decay in optical density. Substantial death in gram-positive bacteria may be explained by autolysis due to rapid loss of energy-generating substrates (19) or sugar-driven phage induction that leads to lysis (20). Because the substrate concentration in Biolog PM plates is proprietary information and thus unknown, we wanted to see if this phenomenon occurs at different concentrations of these substrates. We therefore assayed 11 clinical isolates of *C. difficile* representing four different ribotypes (a molecular classification of closely related strains) for their growth dynamics on minimal media supplemented with either fructose or glucose as the sole carbon source (**Table S1**). We recapitulated the rapid growth followed by rapid death phenomenon at low 20 mM concentrations of fructose and glucose (**Fig. 3A**). Higher concentration of 50 mM sugar did not result in the rapid decay experienced at low concentrations within 24 hours, although strains reached similar carrying capacity on both concentrations (**Fig. 3A**). Next, we estimated growth parameters and their 95% credible intervals on pooled experimental and technical replicates for each unique combination of ribotype, sugar, and substrate concentration (**Fig. 3B**). Growth on fructose exhibited more dramatic decay of OD after reaching carrying capacity. Indeed, death rates were also higher (more negative rates) at low concentrations of fructose than low concentrations of glucose. In addition, differences in death rates between low and high concentrations were statistically significant for three ribotypes on fructose but only for one on glucose (**Fig. 3B**). In summary, low concentrations of simple sugars, such as glucose or fructose, can result in rapid lysis of *C. difficile* after the stationary phase. Because this phenomenon varies by ribotype, we speculate that it may vary based in part on evolutionary adaptations of different lineages.

**Figure 3.**
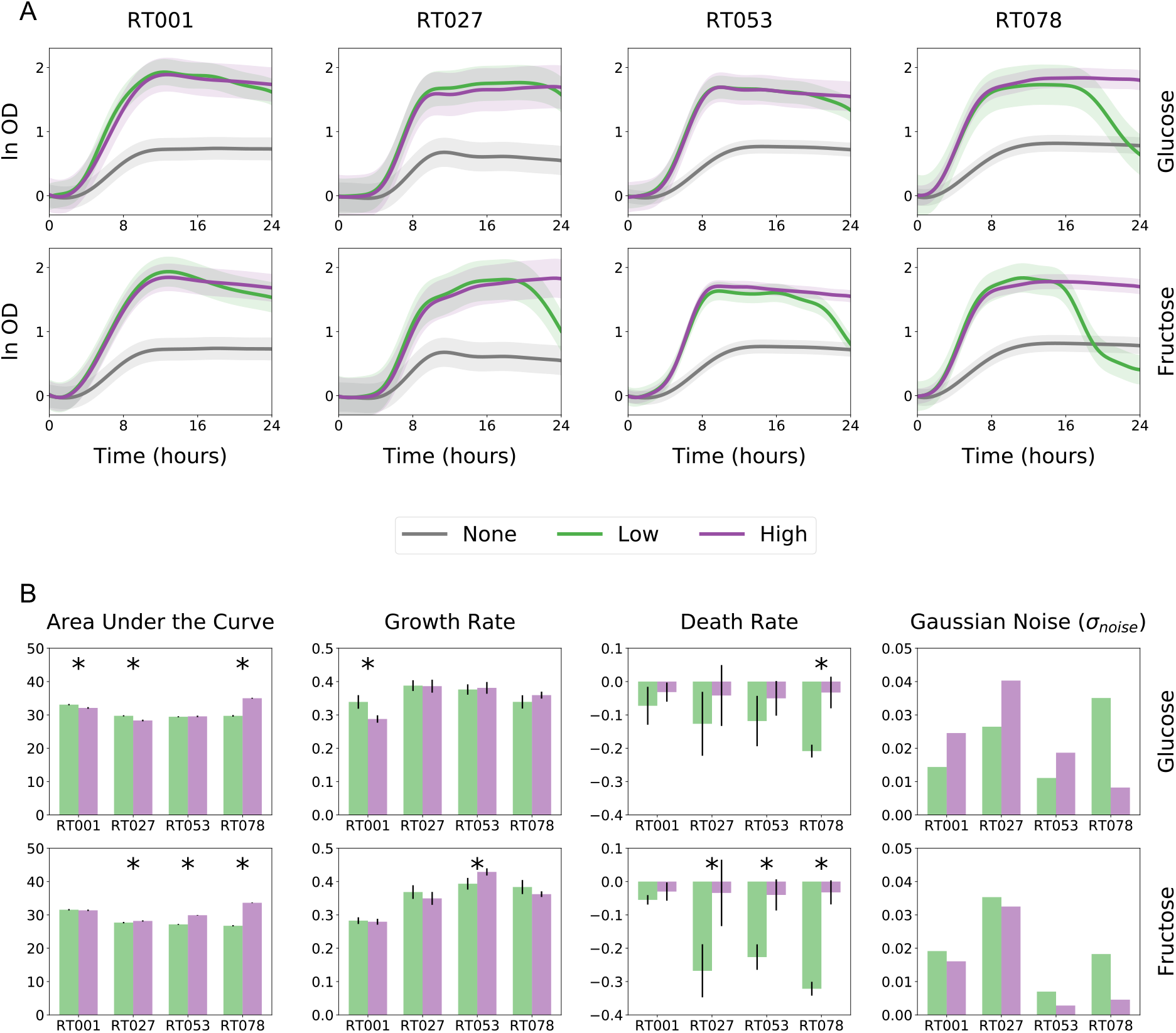
Low concentration of glucose and fructose induce rapid death of microbial cultures in stationary phase. (**A**) AMiGA-predicted growth curves for clinical isolates belonging to four ribotypes (two RT001 isolates, four RT027 isolates, two RT053 isolates, and three RT078 isolates) grown on minimal media supplemented with no (0 mM), low (20 mM), or high (50 mM) concentration of either glucose or fructose. Growth curves for all isolates were measured with three technical replicates. Bold lines indicate the predicted mean of growth and bands indicate the predicted 95% credible intervals including measurement noise. (**B**) Bottom panels summarize differences in growth using model estimates of the area under the curve, exponential growth rate, and stationary death rate. Sampling uncertainty was summarized with the model-estimated Gaussian noise. Error bars indicate the 95% credible interval and asterisks indicate no overlap of credible intervals between low and high conditions.

Ribotypes indeed varied in the precision of their growth dynamics. For example, predicted growth curves for ribotype 027 strains had much wider confidence intervals than curves for ribotype 053 strains (**Fig. 3B**). The difference in confidence is due to difference in experimental variation across biological and technical replicates of each ribotype (**Fig. S5**). Difference in measurement variance is also captured by the model-optimized Gaussian noise term which is used in the estimation of confidence intervals (**Fig. 3B**). Here, noise is modelled as a single time-independent term and optimized for estimating measurement noise across all time points while maximizing model fit. To do so, it may however amplify confidence intervals at time points where measurement variance is actually low. For example, measurement noise is smaller during lag and exponential phases but larger during stationery and death phases. To fine-tune predicted confidence intervals, AMiGA users can opt to empirically estimate measurement noise as previously described (9) which can result in confidence intervals that more closely follow actual measurement noise over time (**Fig. S5**). Still, the time-independent noise term provides useful insight about our experiment. Here, we saw that ribotype 027 has the largest noise term and widest confidence intervals relative to other ribotypes. This may reflect larger phenotypic diversity amongst ribotype 027 isolates or may suggest a need for generating starting cultures in a manner that reduces variability in initial population size and physiology (21).

### Quantifying differential growth across all time points

We initially analyzed experimental conditions by comparing growth parameters using standard univariate statistical tests to detect significant differences attributed to a particular treatment or condition. This approach only detected differences captured by selected growth parameters. AMiGA can instead agnostically test for differential growth due to specific covariates (or conditions) as previously developed and described (10, 22). Briefly, the primary covariate of microbial growth measurements is the independent variable of time. We can extend a GP regression model to include additional categorical covariates that may contribute to differences in growth measurements over time such as nutrient conditions (e.g. carbon substrate) or genotypes (e.g. *C. difficile* ribotype). We can then jointly predict microbial growth on these different conditions and the functional difference in growth between these conditions (**Fig. 4A**, **Table S2**). Functional differences that deviate from zero suggest that different experimental conditions yield different growth dynamics. By summarizing these functional differences into a single metric, we can further compare how sugar concentration contributes to differences in growth based on *C. difficile* ribotype (**Fig. 4B**). In particular, AMiGA computed the squared sum of the functional differences over time, ∥*OD*Δ∥, and confirmed that RT078 isolates experienced the largest growth differences due to sugar concentration and that differences of growth due to fructose concentration are higher than due to glucose concentration (**Fig. 4B**).

**Figure 4.**
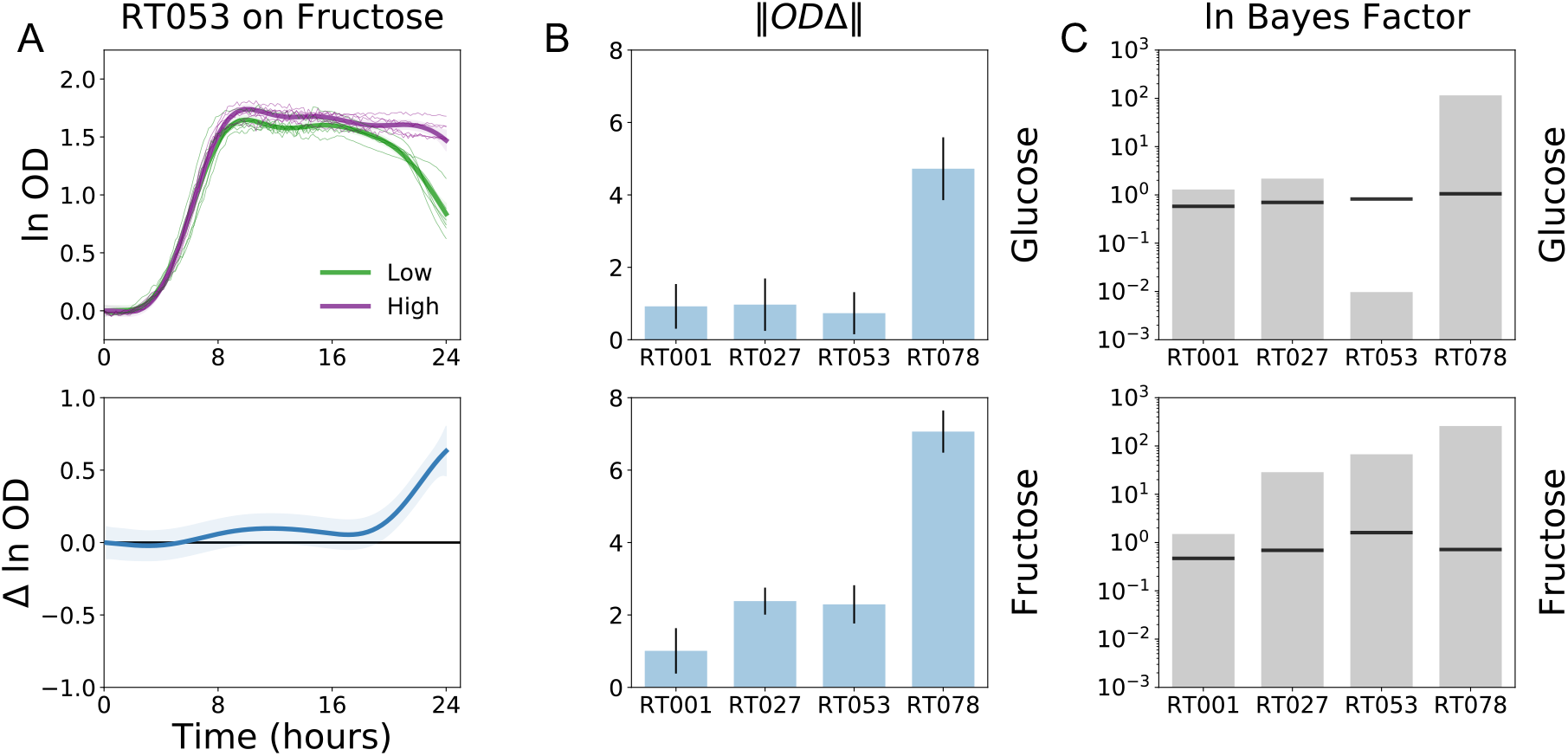
Testing for differences in growth of each ribotype on low versus high concentration of glucose or fructose. Growth of strains belonging to each ribotype on different concentrations of each sugar were modelled jointly. **(A**) Ribotype 053 *C. difficile* exhibited functional differences (Δ ln OD) in growth with small differences in OD during early stationary phase and much larger differences in OD after 18 hours. **(B)** Functional differences between growth on low versus high concentration of each sugar are summarized with the squared sum of functional differences. Error bars indicate 95% confidence intervals. **(C)** Log Bayes Factor scores probabilistically estimated how much model performance improves by including the concertation covariate in addition to time in the GP regression model when comparing growth on different concentrations of sugar for each ribotype. Black horizontal lines indicate the FDR≤10% threshold based on 100 model permutations. Actual log Bayes Factor scores above these thresholds are considered significant.

The functional difference metrics is valuable for rank ordering conditions that impact overall growth dynamics. However, these summary scores are sensitive to the scale or magnitude of the growth curves. If the overall growth of *C. difficile* on both conditions were both amplified by the same factor, so will the functional differences between them. We can instead quantify the effect of an additional covariate on the growth model with less bias from the overall magnitude of growth using log Bayes Factor scores (10). Briefly, growth is modelled on two hypotheses: the null hypothesis assumes that only time explains differences in growth data while alternative hypothesis includes additional covariates of interest in the model. We assess the goodness of fit (or likelihood) of these competing models using the log Bayes Factor score which is the log ratio of the likelihood of the alternative hypothesis to the likelihood of the null hypothesis. Log Bayes Factor scores higher than zero provide more evidence for the alternative hypothesis than the null hypothesis. By permuting the covariate labels, we can also determine the False Discovery Rate (FDR) thresholds and only consider Bayes Factor scores that outperform the threshold as significant. We can broadly recapitulate the above findings using the nonparametric log Bayes Factor score which showed that substrate concentration improves model performance for all conditions except for RT053 isolates grown on glucose (**Fig. 4C**). Importantly, the log Bayes Factor score highlights differences in growth curves that are not captured by significant differences in death rate, such as the differential growth of RT001 and RT027 on glucose as well as the differential growth of RT001 on fructose (**Fig. 3B**). Although significant, these differences are minor in terms of functional differences or effect size as indicated by the sum of predicted functional differences ∥OD_Δ_∥ (**Fig. 4B)**. Overall, differential testing can help users prioritize post-hoc analysis and experimental validation by ranking conditions based on log Bayes Factor scores or squared sum of functional differences.

## DISCUSSION

Recent advances in laboratory automation have dramatically increased the throughput of microbial growth assays. Important aspects of analyzing these data sets include screening for desired phenotypes (23), discovering genotype-phenotype relationships (22, 24), investigating crosstalk of environmental pressures and microbial dynamics (25, 26), and predicting microbial fitness under a variety of conditions (27, 28). Yet, we are continuing to learn how growth dynamics are affected by a variety of technical and experimental factors (21). Scientists are also increasingly able to cultivate nontraditional or non-lab adapted strains and manipulate them under a wide variety of treatments. The study of diverse microbes under widely different applications and growth conditions can thus generate growth modalities that do not follow standard sigmoidal growth and require more nuanced analysis. Here, we contribute a user-friendly software that can further streamline the Analysis of Microbial Growth Assays (AMiGA) with a nonparametric modelling approach.

We showcased our software using multiple examples to highlight several useful features. In particular, AMiGA can infer biologically meaningful parameters either by analyzing individual growth curves or pooling replicates of similar experimental conditions across multiple data sets. The latter approach enables inference with summary statistics for growth parameters, in particular the mean and confidence intervals. It can also probabilistically compare microbial growth under distinct conditions which can aid scientists in ranking experimental conditions for post-hoc validation. We also developed a novel algorithm for the detection and characterization of multi-phasic growth without any underlying assumptions of growth curve shape. Diauxie is a complex behavior characterized by two phases of growth that are often separated by a lag phase (1, 12, 29). There are no consensus formulations for diauxie, and current studies have relied on ad hoc analysis of growth data. Our novel algorithm for diauxie characterization can simplify its analysis and contribute to ongoing efforts for studying its behavior. Finally, emerging research on antibiotic- and phage-microbial interactions can benefit from automated approaches for measuring and quantifying the inhibition of microbial growth or death of microbial culture as we demonstrate here (23, 28). In summary, AMiGA streamlines the analysis and interpretation of growth curve assays in a high-throughput user-friendly manner and can be utilized in a wide variety of microbiology experiments.

## MATERIALS AND METHODS

### Bacterial strains and growth

*C. difficile* strains were mostly clinical isolates obtained from the Michigan Department of Community Health (**Supp. Table 1**) (30). Growth assays were carried out at 37°C in anaerobic atmosphere (5% CO_2_, 5% H_2_, 90% N_2_) using pre-reduced media. Strains were cultured overnight in BHI media (Difco) supplemented with 5% (w/v) yeast extract. *C. difficile* cultures were diluted 1:10 in DMM (31) to a final OD of ~0.05. Cultures were mixed 1:1 with sugar solutions for final concentrations of either 20 mM or 50 mM of glucose or fructose. For Biolog PM assay, overnight growth of CD2015 was sub-cultured 1:5 in fresh BHIS broth and grown to mid-exponential phase (OD ~0.6) then diluted in DMM to a final OD of 0.05. Each well in the pre-reduced Biolog PM plate was inoculated with 100 uL of final cell suspension. In remaining assays, cultures were grown in 200 uL volumes. All cultures were grown for 24 hours and optical density (620 nm) was measured every 10 minutes immediately after 5 seconds of orbital shaking.

### Modelling growth data as a Gaussian Process

Microbial growth is defined as the observations of microbial abundance over time. Mathematical models such as logistic or Gompertz equations can be used to describe a microbial growth curve (32). However, microbial growth does not always follow the standard sigmoidal shape. As a nonparameteric statistical approach, GPs can model microbial growth without making assumptions about the underlying shape or characteristics of its function. GP regression has several advantages over other nonparametric approaches such as spline fitting and polynomial regression. In particular, GP regression is more robust to outliers and technical variation, can inherently pool replicates, infer derivatives, and estimate growth parameters without cross-validation or bootstrapping (9, 10).

A GP is a probability distribution over functions (here, curves) where any finite number of observations of these functions, with independent and identically distributed Gaussian noise, are distributed as a multivariate normal distribution (33):

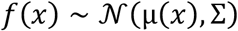

To model microbial growth curves, a GP can be indexed at time such that a microbial growth function is a vector where each entry in the vector specifies a random variable, here, the function value *f(t)* at a particular input *t*. While a Gaussian Process is a distribution over an infinite number of arbitrary functions, we can bias a GP to infer functions that follow certain characteristics. To ensure the inference of smooth microbial growth functions, we specify the priors of a GP to a mean of zero and the kernel or covariance function to a radial basis function (RBF), which is equivalent to a squared exponential, with time-independent Gaussian noise hyperparameter. Accordingly, the full model is specified as

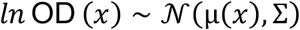

with priors

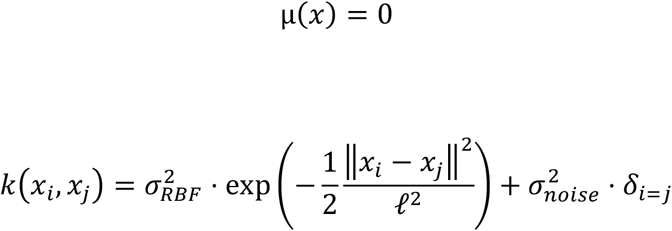

where *i* and *j* are two time points, ∥·∥^2^ is the Euclidean distance, and model hyperparameters include 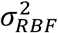, RBF variance, 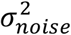, Gaussian noise, and *ℓ*^2^, lengthscale.

In the above definition, the sampling uncertainty is modelled by the time-independent Gaussian noise hyperparameter, 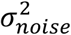. Users can opt to empirically estimate measurement noise and include 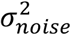 as a time-dependent parameter as previously described (9). In particular, we specify the kernel to be the sum of the RBF kernel and a diagonal matrix of time-dependent noise parameters. These parameters are estimated empirically using the variance across all replicates at each time point, which is then smoothed over time with a Gaussian filter. Unlike model-estimated noise, empirically estimated noise is only used for model optimization and cannot be used for predicting OD at new time points.

### Estimating microbial growth parameters

The GP model parameters are optimized to any given data by maximizing the marginal likelihood through integrating over all possible functions. The optimized hyperparameters can then be used to predict the latent or hidden function and sample new functions from its posterior distribution. Because the derivative of a GP is another GP (33, 34), we can also use GPs to make predictions about the derivatives of the growth curve (i.e. growth rate over time) and sample the posterior of the first and second derivatives of the GP.

AMiGA estimates growth parameters from the optimized model either directly by analyzing the latent function and its derivatives or by sampling many functions from the posterior of the latent function and its derivatives then estimating the growth parameters of each new sample. The latter approach provides a distribution for the estimates of each growth parameter which are then summarized into their means, standard deviations, and confidence intervals.

The growth curve metrics of carrying capacity, *K*, maximum specific growth rate, *μ*_*max*_, and area under the curve, *AUC*, were estimated as previously described (10). Briefly, *K* and *μ*_*max*_ are the maximum a posteriori (MAP) estimates of the *ln* OD (*t*) and the (*d*/*dt*) *ln* OD (*t*) functions. The estimate of *AUC* was calculated as the Riemann sum of the *ln* OD (*t*) function. In particular, *ln* OD (*t*) was predicted at evenly space time points, then linearly transformed with vector of time intervals *a* = Δ*t*, Δ*t*, … such that an approximation of the AUC follows a normal distribution of 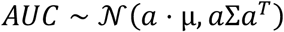.

AMiGA also computes additional growth parameters including death, maximum death rate, adaptation time, lag time, and doubling time. Death is simply the absolute difference between the growth measurements at the final time point and the carrying capacity. The maximum death rate is estimated as the minimum of the negative of the derivative function (*d*/*dt*) *ln* OD (*t*) where *t* is limited to (t_k_, … , T) to capture maximum death rate after reaching carrying capacity at *t*_*K*_. Adaptation time is computed probabilistically as the time at which the 95% credible interval of the growth rate, (*d*/*dt*) *ln* OD (*t*), deviates from zero. Lag time is computed using the classical definition of lag time (32) as the intersection of the tangent line to the (*d*/*dt*) *ln* OD (*t*) function at maximum growth rate and the line parallel to the x-axis or time (9). Doubling time is computed as *ln*(2) times the inverse of the maximum specific growth rate.

### Detecting and characterizing multiple growth phases

AMiGA applies a novel algorithm for detecting diauxic shifts that utilizes first- and second-order derivatives of growth measurements and a customizable heuristic for calling unique growth phases (**Supp. Fig. 1**). AMiGA computes the first- and second-order derivatives of each latent function which correspond to the growth rate and the change in growth rate over time respectively. The second-order derivative indicates inflection points that are defined as (*d*^2^/*dt*^2^) *ln* OD (*t*) = 0. Positive inflection points indicate acceleration of growth while negative inflection points indicate deceleration. These inflection points also correspond to the valleys and peaks in the first-order derivative respectively. Each potential unique growth phase is bounded by two consecutive positive inflection points. AMiGA handles edge measurements by assuming positive inflection points at the first and last time points.

AMiGA then applies a heuristic to determine if each potential growth phase actually indicates a significant change in growth dynamics. In an iterative process, AMiGA compares potential phases based on their total growth. Starting with the phase with the smallest growth, AMiGA compares it to the primary phase with the largest growth. If the smallest phase is associated with growth that exceeds a certain ratio of the growth caused by the primary phase, it is confirmed as a unique growth phase. Otherwise, this non-growth phase is merged to one of its adjacent phases. The non-growth phase is merged to the adjacent phase with the lower activation energy, which is defined as the difference between the maximum growth rate during the non-growth phase and the growth rates at either its left or right boundary (i.e. at its beginning and end). Growth phases are then re-ranked based on their total growth and iteration continues.

The iterative process is completed when the smallest growth phase is confirmed as a unique growth phase or all potential phases have been merged into a single growth curve and thus indicating that no diauxic shifts (or shifts in growth phases) have occurred. Users can alter several parameters of this algorithm including the threshold or proportion that defines a unique growth phase (default is 20% of growth attributed to the primary phase) and whether to compare potential growth phases based on total change in growth measurements, *ln* OD (*t*), or total change in growth rate, (*d*/*dt*) *ln* OD (*t*). Once all growth phases have been detected, AMiGA describes them by their growth parameters in addition to their time boundaries.

### Testing for differential growth

AMiGA applies the framework of Tonner et al. (10) for Bayesian testing of differential growth between two conditions. A Gaussian Process can be extended to model additional covariates or dimensions beyond time where each dimension has a unique independent lengthscale. This is illustrated in vector notation of the covariance as described in Solak et al. (34) where

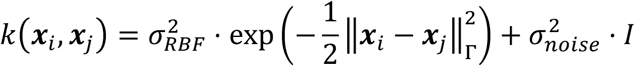

and the norm ∥ ⋅ ∥_r_ is defined as

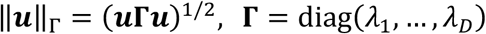

Accordingly, AMiGA tests for differential growth by comparing a null model where the only dimension in input is time, ***x***_*i*_ = {*time*}, with an alternative model where input is multi-dimensional, for example, ***x***_*i*_ = {*time*, *substrate*}, with time as the first covariate and an additional covariate included as a binary variables *x*_*i*,1_ ∈ {0,1}.

Differential growth is quantified with a Bayes Factor score defined as the ratio of the likelihood of the data given the alternative hypothesis or model to the likelihood of the data given the null hypothesis or model.

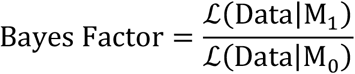

A Bayes Factor score higher than 1 indicates stronger evidence for the alternative model (M_1_) than the null model (M_0_). High scores indicate that the additional covariate improves null model performance and suggest that growth curves are functionally different due to covariate effect. A false discovery rate (FDR) for each Bayes Factor score is calculated by estimating the null Bayes Factor score distribution. In particular, the label of the additional covariate was randomly assigned to each sample without replacement from the original distribution of the covariate in the model input. Then, a null Bays Factor score was calculated for each permutation of the model. The null distribution comprised 100 permutations of the data set and FDR (default is ≤ 10%) was defined as the 90^th^ percentile of the null Bayes Factor distribution.

### Quantifying functional differences between conditions

When testing for differential growth, AMiGA also models the functional difference due to the additional covariate. The functional difference across all time points is defined as the difference in OD predicted for each condition (10) as follow:

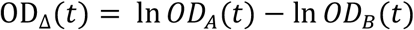

where OD is predicted for conditions A and B jointly. These differences can be summarized into a single metric, ∥OD_Δ_∥, which is the squared sum of predicted OD differences and represents the magnitude of functional difference between conditions (22).

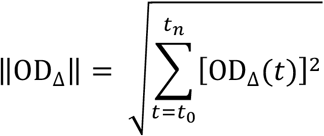

The uncertainty of the squared sum of functional differences is estimated by sampling 100 times from the posterior distribution of the functional difference, computing the squared sum of functional differences for each sample, then reporting the mean and 95% confidence interval of the distribution of squared sums.

### Estimating confidence intervals

AMiGA has options for exporting preprocessed and postprocessed data, including input and output of the GP model, predicted growth curves either in log or linear scale, predicted mean and covariance for both growth and its first-order derivative, predicted Gaussian noise, 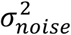, as well as sampling uncertainty (i.e. measurement variance) if estimated empirically. Predicted means and covariances can then be used to compute 95% credible intervals as follow:

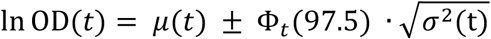

Here, *μ*(*t*) and *σ*^2^(t) are the mean and variance of the latent function respectively, and Φ_t_ is the inverse of the cumulative distribution function for the standard normal. The total variance for predicted OD at each time point is computed as the sum of the variance of the latent function, *σ*^2^(t), and variance due to Gaussian noise, 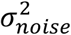. Users can opt to plot confidence intervals and/or compute functional differences without measurement noise in differential growth analysis.

### Data processing

AMiGA can ignore the first *n* time points because plate reader measurements often exhibit the highest variation in OD at the first few time points. Here, we only eliminated the first measurement of each growth curve. OD values were transformed with natural logarithm. The OD value at the first measurement was then estimated with polynomial regression of degree of five on the first five time points as previously described (10). This estimated OD measurement was then subtracted from all consecutive measurements in a curve. In differential testing, growth curves belonging to group-specific control samples (e.g. no-carbon control in Biolog or water control in other assays) can be subtracted from target growth curves. In our analysis, growth curves were not normalized to control samples. Growth curves can be visualized in linear scale by exponentiating the model estimate of the latent function and assuming starting OD of 1. In addition, area under the curve, carrying capacity, and death values can be estimated either in log or linear scale based on user preferences.

### Software implementation and data availability

AMiGA is written in Python 3 (Python Software Foundation, https://www.python.org). It utilizes GPy for Gaussian Process regression (36); Pandas (37), NumPy (38), and SciPy (39) for data manipulation and scientific computing; and MatPlotLib (40) and Seaborn (41) for data visualization. Code for AMiGA and growth data analyzed in this article are hosted online (https://github.com/firasmidani/amiga) with detailed documentation and tutorials.

## Acknowledgements

We thank Dr. Lei Pan (Baylor College of Medicine) for sharing Biolog PM1 data for CD2015, a ribotype 027 *C. difficile* isolate.

This work was supported by National Institutes of Health (NIH) grants to R.A.B (U01AI124290 and R01AI123278). F.S.M was supported by an NIH institutional training grant (T32DK007664). R.A.B. receives unrestricted research support from BioGaia AB, consults for Takeda and Probiotech, serves on the scientific advisory board of Tenza, and is a cofounder of Mikrovia.

## SUPPLEMENTAL TABLES

**Supplemental Table 1.** Identity and types of strains used in this study. Strains are clinical isolates collected and provided by the Department of Michigan Community Health, except for M120 which was provided by Ed Kuijper (Leiden University Medical Center, Leiden, The Netherlands). NA indicates not available.

| Strain | Toxintype | PFGE type<br>(NAP status) | Ribotype |
| --- | --- | --- | --- |
| CD2015 | III | MI-NAP1 | RT027 |
| CD4015 | III | MI-NAP1 | RT027 |
| CD4001 | III | MI-NA13 | RT027 |
| CD4010 | III | MI-UN13 | RT027 |
| CD1015 | V | MI-NAP7 | RT078 |
| M120 | V | NA | RT078 |
| CD2001 | V | MI-NAP8 | RT078 |
| CD3014 | 0 | MI-NAP2 | RT001 |
| CD4011 | 0 | MI-NAP2 | RT001 |
| CD1007 | 0 | MI-NAP3 | RT053 |
| CD2048 | 0 | MI-NAP3 | RT053 |
| CD2058 | NA | MI-UN5 | NA |

**Supplemental Table 2.**
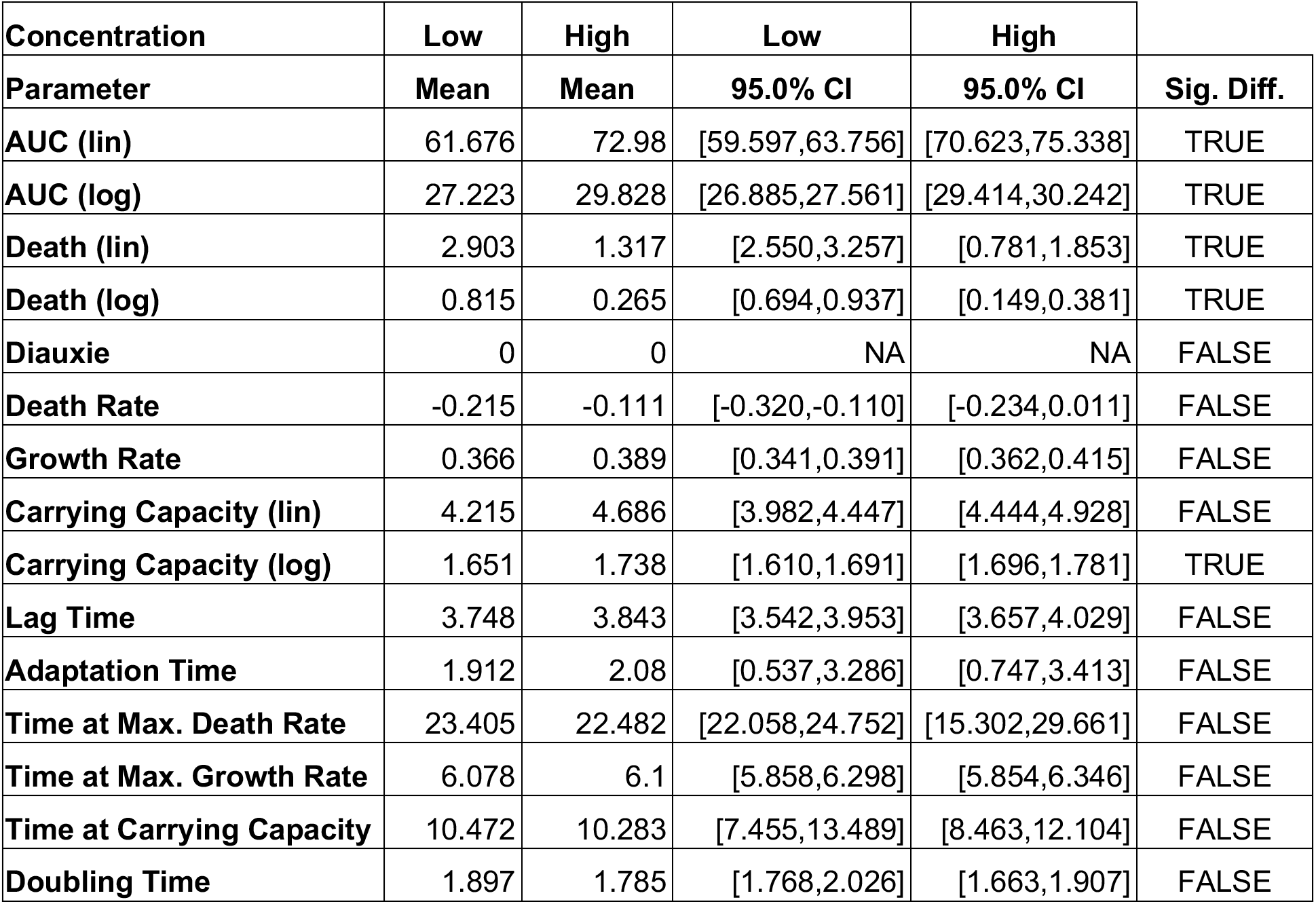
AMiGA-generated comparison for a joint model of growth curves of ribotype 053 *C. difficile* clinical isolate grown on either low (20 mM) or high (50 mM) fructose concentration. Table reports the estimated mean and 95% confidence interval for growth parameters. The index column describes condition or growth parameter. The “Sig. Diff.” column describes whether there is a significant difference between conditions based on whether the 95% confidence intervals overlap or not. For diauxie, “1” and “0” indicate true or false respectively. “NA” indicates not available. “Lin” and “log” indicate whether a growth parameter was calculated using linear or logarithm-scaled optical density values. Measurements based on a linear scale assume a starting OD of one for the first time point. Linear measurements therefore estimate fold-changes over time.

## SUPPLEMENTAL FIGURES

**Supplemental Figure 1.**
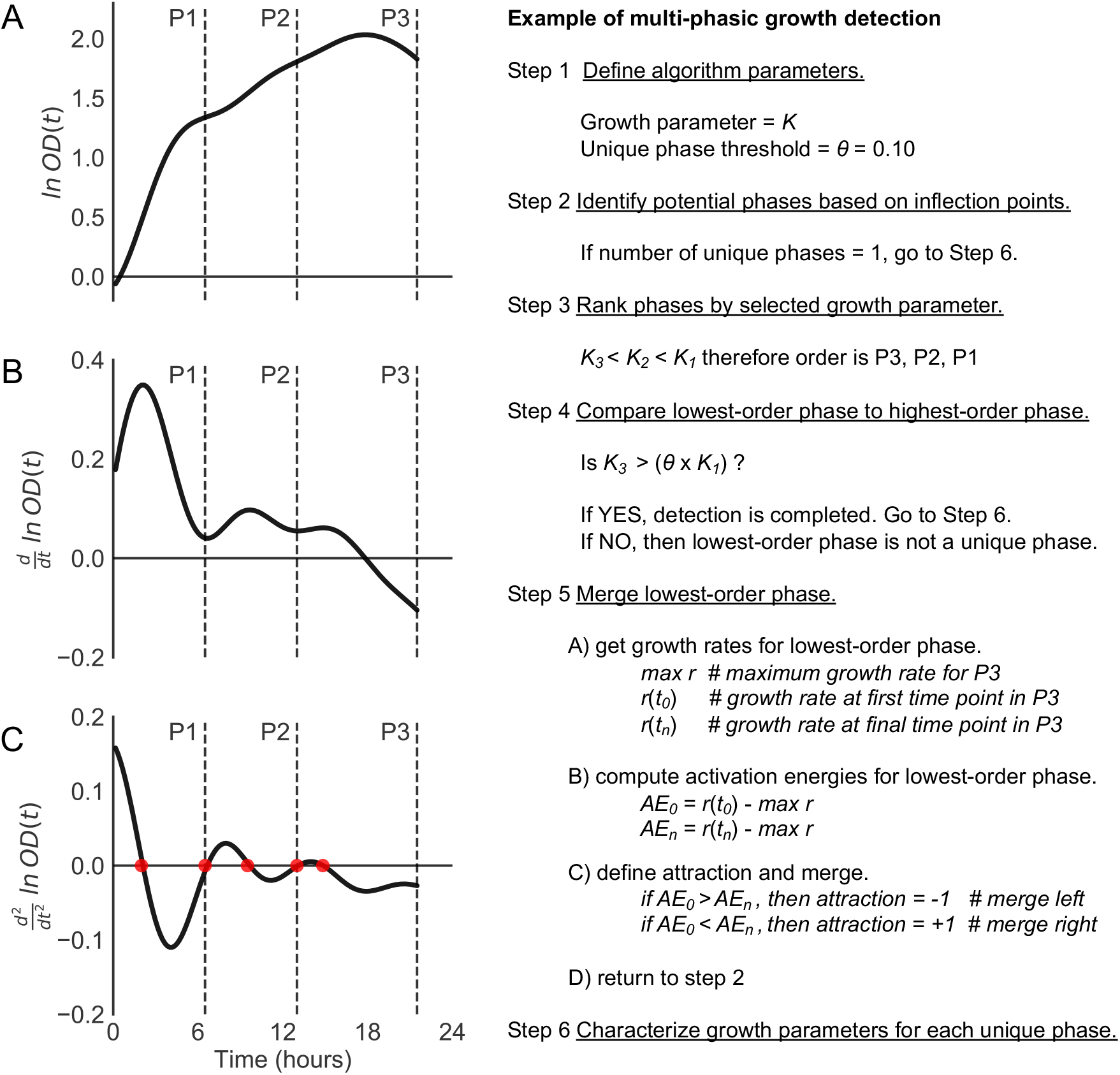
Illustration of diauxic shift detection algorithm using first- and second-order derivatives of growth. Here, we analyze the growth of CD2015, a ribotype 027 *C. difficile* isolate, on sorbitol in Biolog PM1 plates. AMiGA determined all potential growth phases based on inflection points which are defined as the time points at which the second-order derivative equals zero (red points in **C**). Each potential growth phase is bounded by two positive inflection points which correspond to valleys in the first-order derivative (**B**, Step 2). Potential growth phases are then ranked based on carrying capacity (*K*) or change in OD during their time intervals (**A**, Step 3). In this example, the growth phases are ranked in ascending order as P3, P2, P1. The smallest growth phase (P3) does not meet the threshold of producing 20% or higher growth relative to the primary growth phase (P1) (Step 4). The smallest phase (P3) is then merged with the neighboring growth phase (P2) because the maximum growth rate (*r*) for the smallest phase around 16 hours is closer to the growth rate at its left bound around 13 hours (*t*_*0*_) than to its final growth rate at its right bound at 21.7 hours (*t*_*n*_) (**B**, Step 5). The newly defined growth phase that spans both P2 and P3 has a carrying capacity that is at least 20% of the carrying capacity for the largest phase (K_1_) and therefore considered a unique growth phase. In summary, ribotype 027 clinical isolate grown on minimal media supplemented with sorbitol undergoes a diauxic shift with the second growth phase beginning at around 6.7 hours.

**Supplemental Figure 2.**
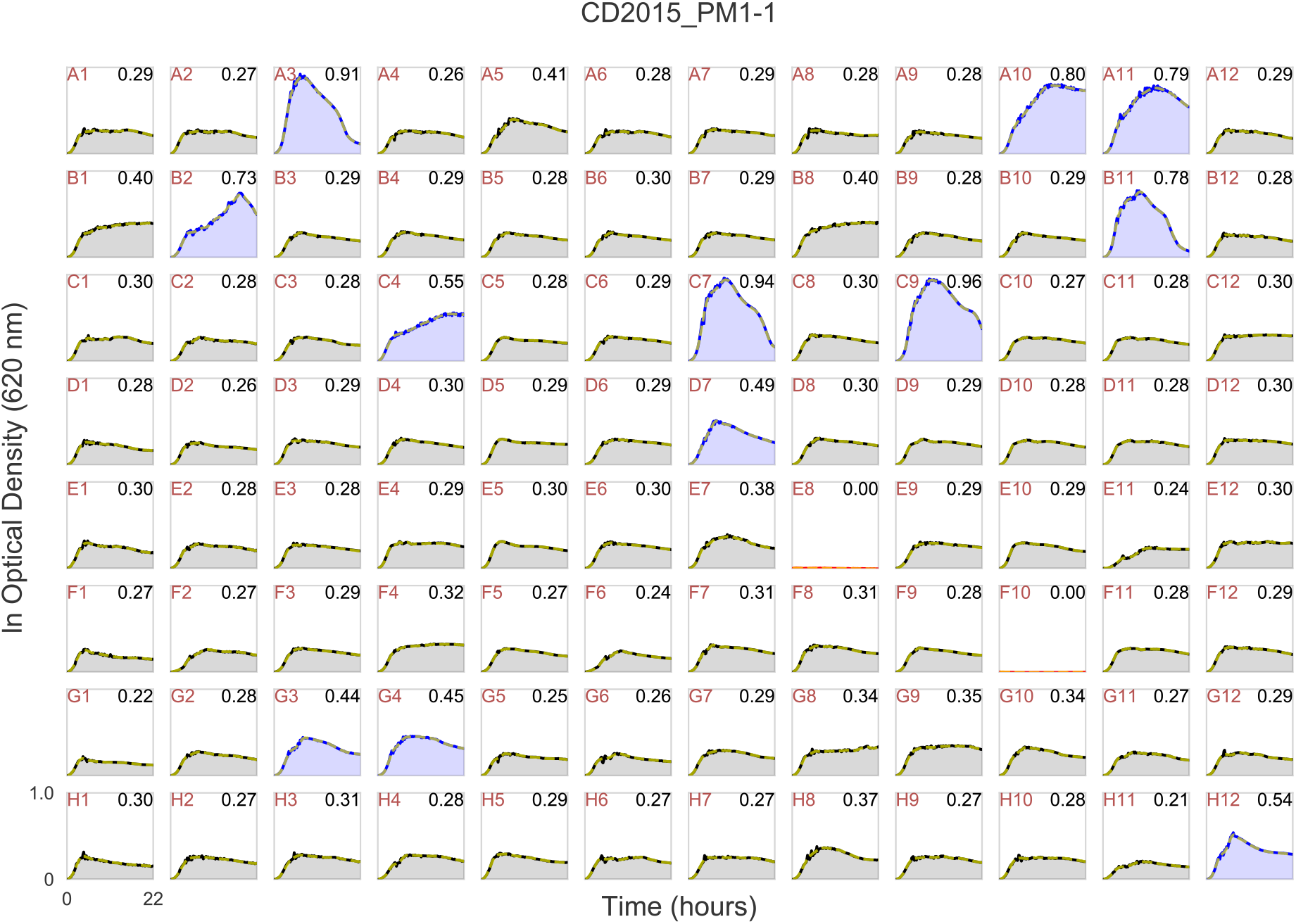
AMiGA visualization of growth curve analyses. AMiGA plotted 96-well grid of growth curves for CD2015, a ribotype 027 *C. difficile* isolate, on a Biolog PM1 plate and highlighted wells where significant growth (blue) or death (red) is detected. Bold lines indicate the actual growth data. Dashed yellow lines indicate the model-predicted growth curves. Text on the top left and top right of each subplot indicate Well ID and the maximum ln OD of each growth curve respectively.

**Supplemental Figure 3.**
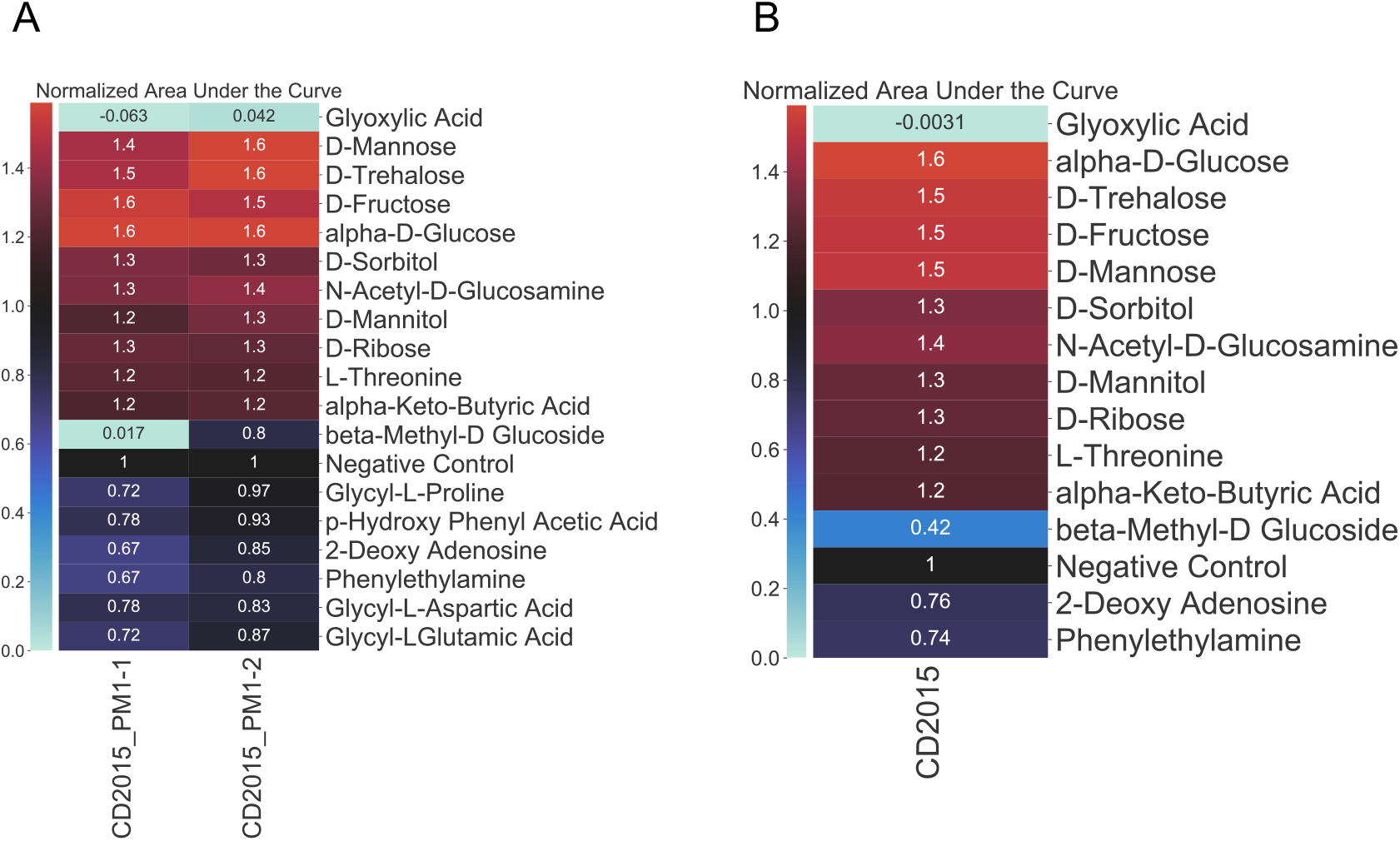
AMiGA summary of Biolog results with heatmaps. Here, we summarized the estimates for normalized area under the curves (AUCs) for two technical replicates of CD2015, a ribotype 027 *C. difficile* clinical isolate, grown on Biolog PM1 plate. (**A**) AMiGA-generated heatmap of normalized AUCs for each technical replicate modeled individually. (**B**) AMiGA-generated heatmap of normalized AUCs for technical replicates modeled jointly. Heatmaps were reduced to substrates that supported a normalized AUC of at least 1.2, equal to 1, or less than 0.8 in either of the technical replicates or in the pooled model. Labels on the x-axis indicate unique plate IDs. In each heatmap, substrates were clustered based on Euclidean distances. In general, heatmaps visually summarized the carbon substrates that support (red) or inhibit (blue) the growth of CD2015. In addition, they demonstrated significant technical variation between the two replicates mostly for the normalized AUCs on substrates that inhibit growth and especially for beta-Methyl-D Glucoside. Most of these variations can be attributed to overall better growth of the first replicate (CD2015_PM1-1) on minimal media without any carbon sources. In summary, the heatmap function aids users in evaluating the quality of their data and model estimates.

**Supplemental Figure 4.**
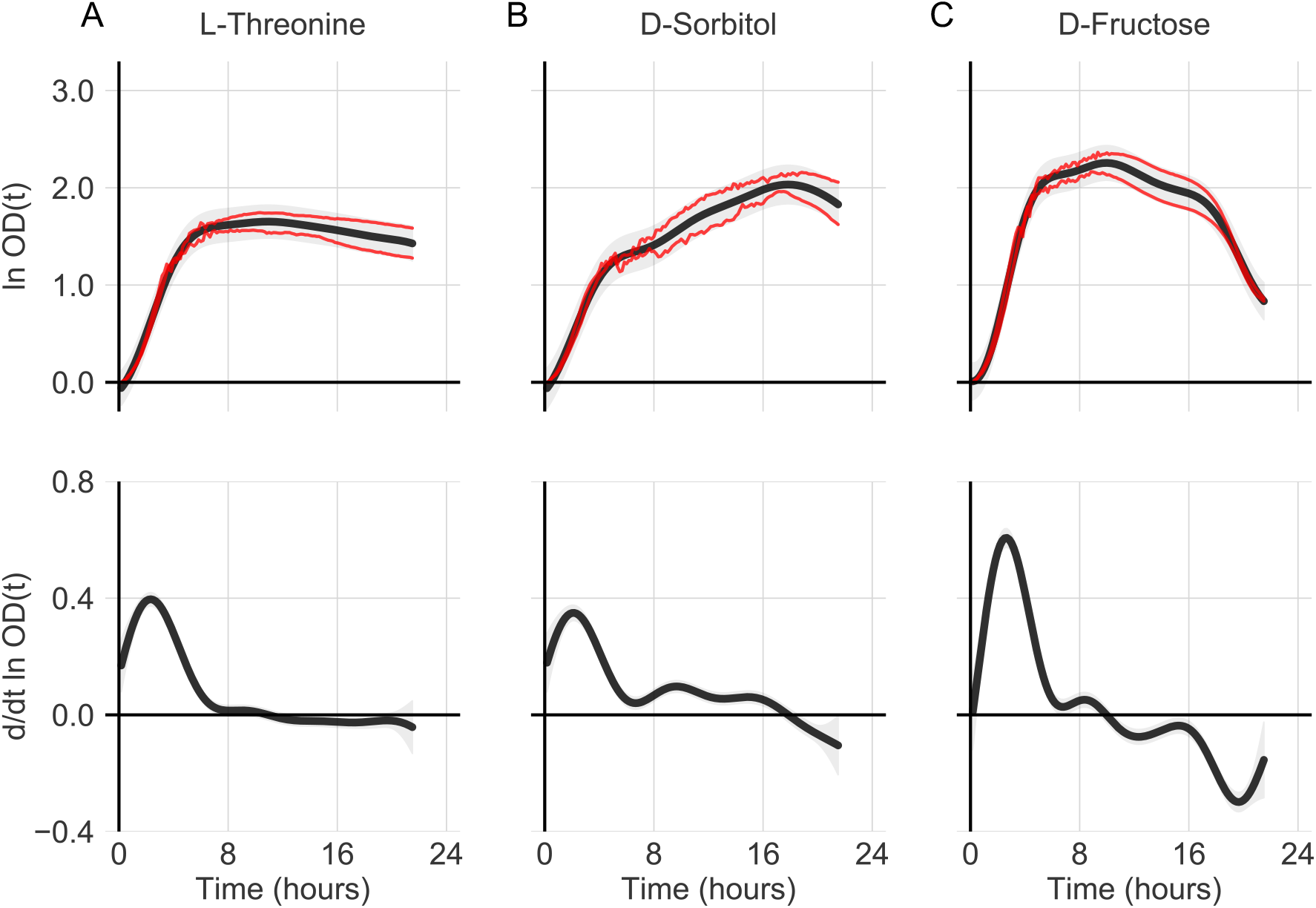
AMiGA can fit growth curves without any underlying assumptions about their shapes. AMiGA predicted growth measurements and growth rates over time for CD2015, a ribotype 027 *C. difficile* isolate, grown in Biolog PM1 plate on minimal media and supplemented with single carbon sources. Bold black lines indicate the mean estimate of the growth function and gray bands indicate the 95% confidence interval including sampling uncertainty (i.e. measurement noise). Red lines plot the actual raw measurement of duplicate growth curves for each condition. The classical logistic growth curve typical of standard microbial cultures is demonstrated by growth of the isolate on L-Threonine (**A**). Exponential growth followed by rapid decay of OD is demonstrated by growth on D-Fructose (**C**). Growth on D-Sorbitol demonstrates a common growth modality for *C. difficile* where OD initially increases exponentially but then increases at lower growth rate for a prolonged period of time, prolonged positive growth rate that slowly decreases over time (**B**).

**Supplemental Figure 5.**
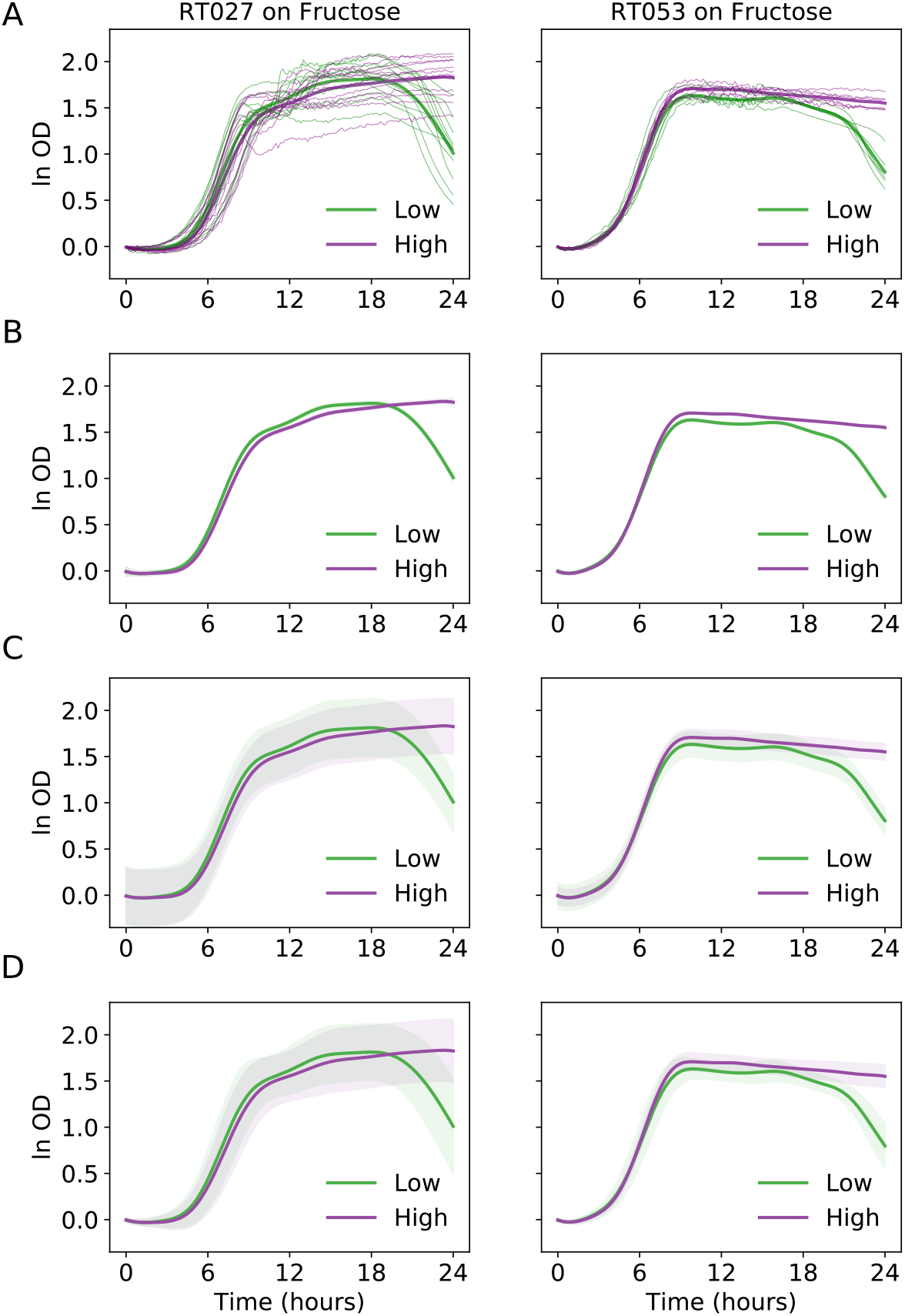
Predicted confidence intervals based on different model manipulations. Growth of four ribotype 027 isolates or two ribotype 053 isolates on different concentrations of fructose were modelled jointly with AMiGA. All strain, substrate, and concentration combinations were repeated in three technical replicates. For all plots, bold lines indicate the predicted mean of growth functions. (**A**) Thin lines show the individual replicate growth curves for each condition. There are 12 replicates for ribotype 027 and 6 replicates for ribotype 053 on each fructose concentration. Ribotype 027 has larger measurement variance or noise than ribotype 053 on both concentrations of fructose. (**B**) Bands, which are barely visible, show the 95% confidence intervals for the growth functions based only on the predicted covariance without the Gaussian noise hyperparameter. (**C**) Bands show the 95% confidence intervals for the growth functions based on the predicted covariance including the gaussian noise hyperparameter. (**D**) Bands show the 95% confidence intervals based on a GP regression model where measurement noise is empirically estimated and included as fixed parameters in the model.

